# Motor Learning Without Moving: Proprioceptive and Predictive Hand Localization After Passive Visuoproprioceptive Discrepancy Training

**DOI:** 10.1101/384941

**Authors:** Ahmed A. Mostafa, Bernard Marius ’t Hart, Denise Y.P. Henriques

## Abstract

An accurate estimate of limb position is necessary for movement planning, before and after motor learning. Where we localize our unseen hand after a reach depends on felt hand position, or proprioception, but in studies and theories on motor adaptation this is quite often neglected in favour of predicted sensory consequences based on efference copies of motor commands. Both sources of information should contribute, so here we set out to further investigate how much of hand localization depends on proprioception and how much on predicted sensory consequences. We use a training paradigm combining robot controlled hand movements with rotated visual feedback that eliminates the possibility to update predicted sensory consequences (‘exposure training’), but still recalibrates proprioception, as well as a classic training paradigm with self-generated movements in another set of participants. After each kind of training we measure participants’ hand location estimates based on both efference-based predictions and afferent proprioceptive signals with self-generated hand movements (‘active localization’) as well as based on proprioception only with robot-generated movements (‘passive localization’). In the exposure training group, we find indistinguishable shifts in passive and active hand localization, but after classic training, active localization shifts more than passive, indicating a contribution from updated predicted sensory consequences. Both changes in open-loop reaches and hand localization are only slightly smaller after exposure training as compared to after classic training, confirming that proprioception plays a large role in estimating limb position and in planning movements, even after adaptation. (data: https://doi.org/10.17605/osf.io/zfdth, preprint: https://doi.org/10.1101/384941)

## Introduction

Sensory information is central to how we control all our movements. Our brain is even thought to use predicted sensory consequences derived from efference copies of motor commands for motor control [1]. When training with rotated visual feedback of the hand, we update these predictions [2], and these updated predictions contribute to motor control. Additionally, such training leads to a recalibration of our sense of felt hand position based on afferent signals (that we call “proprioception” here) to be more aligned with the distorted visual feedback [3]. Both of these changes are measured by asking people to localize where their unseen hand is – before and after training [4–8]. While our lab finds that proprioception accounts for a substantial portion of the change in hand localization, and should hence contribute to motor adaptation [9], it is far from clear how much each process contributes or how to tease them apart. Here we use training with robot-generated reaches, that prevents efference-based predictions and eliminates differences between actual and expected visual feedback and hence prevents updating predicted sensory consequences. We then compare hand localization data from this experiment to hand localization in a classic training paradigm with self-generated movements, from an – otherwise identical – earlier study [9], in an attempt to isolate the contribution of proprioception to hand localization prior to and after adaptation.

The predicted sensory consequences of movements likely play several important roles in motor learning and control. Predicted sensory consequences allow us to correct our movements before sensory error signals are available, they can be used to select movements that best achieve our goals and they may inform us on the location of our limbs. Hence measuring predicted sensory consequences is valuable in movement research. In motor adaptation, the actual sensory outcome is systematically different from the expected outcome, so that participants update their predictions on the outcome of the trained movements. In two previous visuomotor adaptation experiments, people are asked to make a movement and then indicate the location of, or “localize,” their unseen, right hand, before and after training with rotated visual feedback [5, 7]. Since there is no visual information available to the participants, the predicted sensory consequences of the movement should be used in localizing the unseen hand. Both studies find a significant shift in hand localization, providing evidence that predicted sensory consequences are indeed updated as a result of visuomotor rotation adaptation.

However, there is another explanation for shifts in hand localization after visuomotor adaptation. Our lab, and others, have shown that our sense of where we feel our hand to be, proprioception, is also reliably recalibrated after visuomotor adaptation [3, 9–19], and a comparable proprioceptive change is induced with force-field adaptation [20, 21]. Just like efference-based predictions, proprioception also informs us on the location of our limbs. To investigate recalibrated proprioception we on occasion use a task that is very similar to hand localization [3, 6, 22, 23]. Although proprioceptive recalibration is largely ignored as an explanation for changes in hand localization, we and others have shown that it accounts for a substantial part of the changes in localization, along with, but separate from, updates in predicted sensory consequences [9, 19]. Nevertheless, it is far from clear how much proprioception and prediction each contribute to hand localization.

Here we intend to further examine the contribution of proprioception to hand localization, relative to the contribution of predicted sensory consequences. To do this, we use a training paradigm with robot-generated movements, which we refer to as ‘exposure training’, where the cursor always goes directly to the target [19, 24, 25]. This means there is no efference copy available and no visuomotor error-signal, both of which are required to update predicted sensory consequences. That is, exposure training to visuoproprioceptive discrepancy should not lead to updates of predicted sensory consequences. However, the discrepancy between vision and proprioception does drive proprioceptive recalibration. Thus, changes in any kind of hand localization after this type of training should be due to proprioceptive recalibration only. We use the same experimental protocol as before [9], where classic visuomotor training should have changed predicted sensory consequences along with proprioception. This allows comparing shifts in hand localization between the two different types of training, and a better assessment of the contributions of predicted sensory consequences and proprioception to hand localization.

Primarily, we will compare the magnitude of shifts in passive and active hand localization after visuoproprioceptive discrepancy exposure and classic visuomotor training. This is because after exposure training only proprioception should be recalibrated, and since we think this contributes to active and passive hand localization equally, there should be no difference between active and passive localization shifts after exposure training. After classic training, on the other hand, proprioception is recalibrated too but in addition, predicted sensory consequences should also be updated, which only contributes to active localization, so that active and passive localization shifts should be different after classic training. For a secondary analysis we will compare the magnitude of reach aftereffects between exposure and classic training. As exposure training should not lead to updates of internal models, exposure training should lead to lower reach aftereffects than classic training. However, if there are any reach aftereffects at all after exposure training this means that afferent-based localization shifts drive motor changes too, and here we can see to what degree that is the case.

## Methods

We set out to test the relative contributions of proprioception and efference-based prediction to hand localization. We used exposure training, with robot-generated reaches to prevent updates of predicted sensory consequences, but still elicit proprioceptive recalibration, and compared this with a classic training paradigm that should change both predicted sensory consequences and proprioception. We then tested this in both active localization, that should use both sources – updated or not – and in passive localization, that should rely only on proprioception, which should be recalibrated equally by both training paradigms.

### Participants

Twenty-five right-handed participants were recruited for the exposure training group in this study. One participant was excluded for not following task instructions, and three were excluded for low performance on a task that ensures attention during the exposure training. All analyses presented here pertain to the remaining twenty-one participants (mean age: 20.1±2.3, age range: 18-25, 13 females), but the data of the three low-performing participants was included in the online dataset [26] (https://osf.io/zfdth). Participants provided prior, written informed consent as approved by the York Human Participants Review Committee (#2014-240). All had normal or corrected-to-normal vision, and received credit toward an undergraduate psychology course. Participants were screened verbally, and all reported being right handed and not having any history of visual, neurological, and/or motor dysfunction. Published data from 21 further participants (mean age: 24, age range: 18-38, 11 females) that completed classic visuomotor training was used for comparison [9].

### Setup

Participants sat in a height-adjustable chair to ensure that they could easily see and reach all targets. To ensure no visual information on right hand position was available to localize the right hand, the room lights were dimmed and the participants’ view of their right hand was blocked by a horizontal surface, as well as a dark cloth draped between a touch screen and participants’ right shoulders. During all tasks, they held the vertical handle on a two-joint robot manipulandum (Interactive Motion Technologies Inc., Cambridge, MA) with their right hand so that their thumb rested on top of the handle. A horizontal touch screen, reflective surface and monitor were mounted 5, 18 and 29 cm above the handle, respectively. Stimuli were presented using the monitor (Samsung 510 N, refresh rate 60 Hz), which was mounted facing downward, 11 cm above the reflective surface, such that images displayed on the monitor and viewed in the reflective surface appeared to lie in the horizontal plane where the right hand was moving. The reflective surface was mounted horizontally 18 cm above the robot manipulandum. The touch screen was mounted 13 cm underneath the reflective surface (5 cm above the manipulandum), so that subjects could indicate the location of the unseen right hand (specifically the thumb on top of the handle) with their left hand. The touch screen always blocked vision of the right hand, but the left hand was lit up with a small spot light (only in localization tasks) in order to be visible through the reflective surface for accurate touch screen responses. For a schematic illustration of the setup, see Fig 1a.

**Fig 1.**
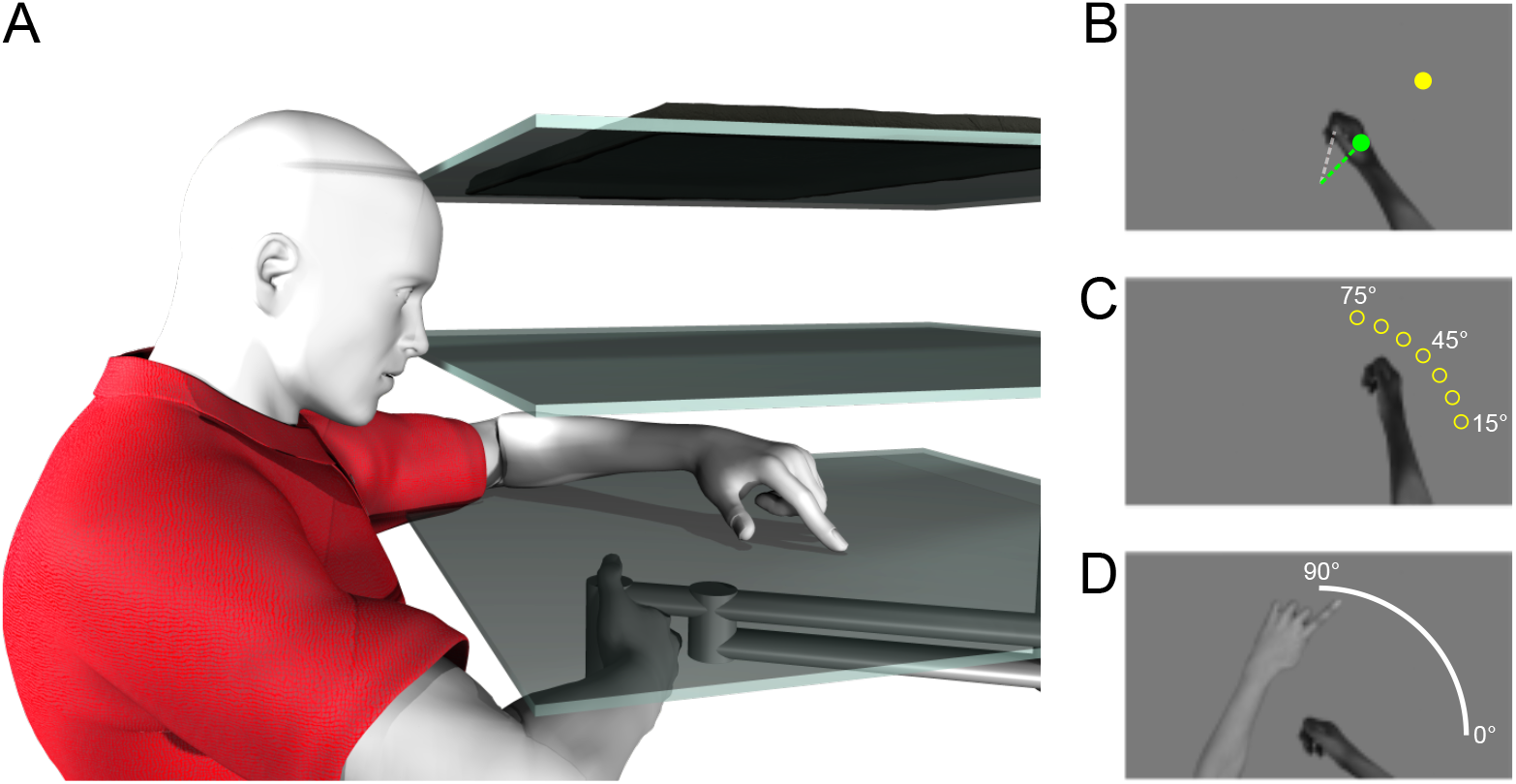
Setup and tasks. **a)** The participants held on to the handle of a robotic manipulandum with their unseen right hand. Visual feedback on hand position was provided through a mirror (*middle surface*) half-way between the hand and the monitor (top *surface*). A touchscreen located just above the hand was used to collect responses for the localization tasks (*bottom surface*). **b)** Training task. The target, shown as a yellow disc, is located 10 cm away from the home position at 45°. In the rotated training tasks, the cursor (shown here as a green circle) represented the hand position rotated 30° relative to the home position. **c)** No-cursor reach task. Targets were located 10 cm away from the home position at 15°, 25°, 35°, 45°, 55°, 65°, and 75°, shown by the yellow circles here (only one was shown on each trial). While reaching to one of these targets, no visual feedback on hand position was provided. **d)** Localization task. The participants’ unseen, right hands moved out and back, and subsequently participants indicated the direction of the hand movement by indicating a location on an arc using a touch screen with their visible left index finger.

### Procedure

The first part of the experiment used training with a cursor aligned with the hand and the second part had training with a cursor rotated around the start position (Fig 1b; Fig 2a). During the training with rotated feedback, the cursor was gradually rotated 30° clockwise. This introduced a discrepancy between the actual, or felt, hand position and the visual feedback, that should evoke proprioceptive recalibration. However, the movements were robot generated, and always brought the cursor straight to the target, so that there were no motor commands generated, and no predicted sensory consequences based on this outgoing command. In the absence of predicted sensory consequences and with perfect, error-free reach performance, there were no sensory prediction errors to drive classic visuomotor adaptation either [27]. After both types of training, participants did two kinds of hand localization tasks, to test the effect of training on proprioceptive and predictive hand estimates, as well as open-loop reaches as a classic measure of implicit adaptation.

**Fig 2.**
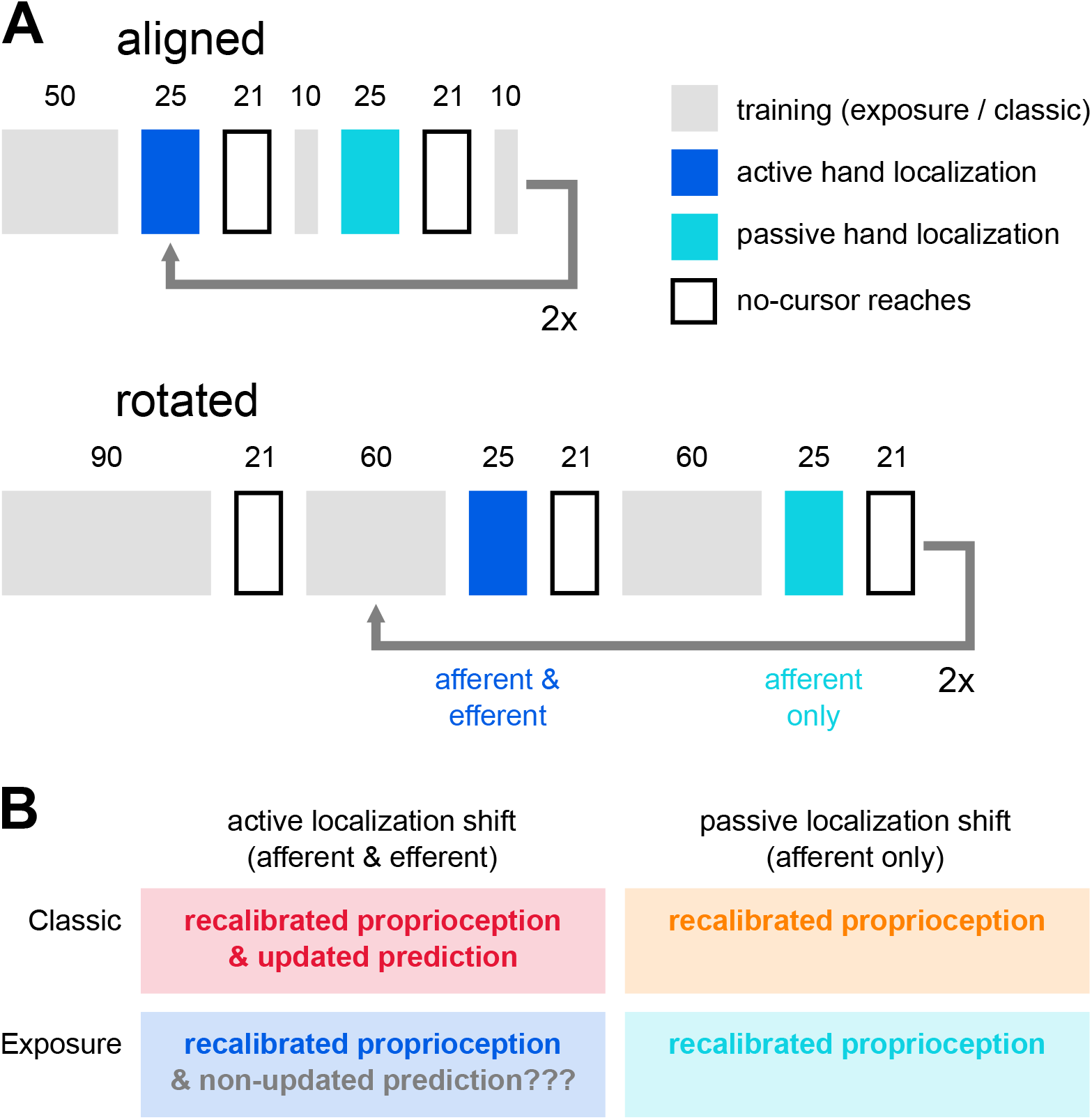
Experimental design. **a)** Task order. All participants performed active and passive localization as well as no-cursor reaching tasks in both a first, aligned session and a second, rotated session. There was an extra no-cursor reach block in the rotated session, to assess decay. The number of trials (numbers above the rectangles) in the no-cursor reaching tasks and both types of localization were the same in the aligned and rotated sessions, but while aligned training was kept to a minimum, rotated training was increased to ensure sufficient adaptation. The exposure training group did passive and active localization twice, but the classic training group did different localization tasks in the second half of each session, the data of which is not used here. **b)** Our main question was about the signals contributing to hand localization in the four different kinds of localization. According to our hypothesis, updated predicted sensory consequences should have only contributed to active hand localization after classic training.

### Exposure training

In what we called ‘exposure training’ the participants did not move their hand toward the target, but the robot did. In this task (Fig 2a, gray boxes), the right hand was represented by a cursor (green disk, 1 cm in diameter, Fig 1b) located directly above the participant’s thumb. The robot moved the participant’s unseen right hand (and the cursor) along a direct path, toward a visual yellow target disk (1 cm in diameter, Fig 1b). The velocity profile was bell-shaped, with a peak velocity of ^~^44 cm/s after ^~^300 ms. The hand reached the target after ^~^600 ms and was then kept in place for an additional ^~^400 ms. The hand was then placed back at the home position using an identical movement. The home position was located approximately 20 cm in front of participants and the visual target located 10 cm from the home position at 45° (Fig 1b). In order to make sure participants were paying attention to the cursor, the cursor was switched off for 2 screen refreshes (^~^33.3ms) on 50% of the trials at a random distance between 4 and 9 cm from the home position and participants were asked to report this using a button press with the left hand. Performance on this task was used to screen participants.

During the first half of the experiment, the “aligned” session, the cursor and hand path were aligned during exposure training. In the second part of the experiment, the “rotated” session, the visuoproprioceptive discrepancy was introduced. The same visual training target at 45° was used, and the cursor kept moving straight to this target. However, the robot-generated hand path gradually rotated 30° CCW (Fig 1b) with respect to the cursor and target, in increments of 0.75°/trial, to prevent awareness of the rotation, so that the full rotation was reached after 40 trials. This mimicked error-free responses to a gradual visuomotor rotation of 30° CW. The initial training consisted of 50 trials in the aligned part and 90 in the rotated part. In between open-loop reach tasks and localization tasks (Fig 2, outlined and blue boxes) extra training tasks were done, each of which consisted of 10 trials in the aligned part of the experiment and 60 trials in the rotated part (Fig 2, gray boxes).

### No-cursor reaching

The trials in no-cursor reaching (Fig 1c; Fig 2, outlined boxes) served to elicit open-loop reaching as a standard measure of motor adaptation. On each of these trials participants were asked to reach with their unseen right hand to one of 7 visual targets, without any visual feedback of hand position. The targets were 10 cm from the home position, located radially at: 15°, 25°, 35°, 45°, 55°, 65°, and 75° (Fig 1c). A trial started with the robot handle at the home position and, after 500 ms, the home position disappeared and the target appeared, cuing the participants to reach for the target. Once the participants thought they had reached the target they held their position (criterion: 4 cm or further from the home position, velocity under 0.5 cm/s for 250 ms), and subsequently the target disappeared, while the home position re-appeared, cuing participants to move back to the home position along a straight, constrained path, to begin the next trial. The path was constrained so that if participants tried to move outside of the path, a resistant force, with a stiffness of 2 N/(mm/s) and a viscous damping of 5 N/(mm/s), was generated perpendicular to the path. In every iteration of the no-cursor reach task, each target was reached for three times, for a total of 21 reaches in pseudo-random order. The no-cursor reaching task was performed four times in the aligned part of the experiment and five times in the rotated part of the experiment. The end-points of no-cursor reaches, in particular the angles relative to the start position and their deviations from the angles of the targets were analyzed, as these typically change in classic visuomotor adaptation.

### Localization

In this task (Fig 1d; Fig 2, blue boxes) participants indicated where they thought their unseen right hand was after a movement. First, an arc appeared, spanning from 0° to 90° and located 10 cm away from the home position and the participants’ unseen, right hand moved out from the home position in a direction towards a point on the arc. The hand was stopped by the robot at 10 cm from the home position and then, to prevent online proprioceptive signals from overriding the predictive signals [5, 9], the hand was moved back to the home position using the same kind of constrained path as used for the return movements in the no-cursor task. Participants indicated with the index finger of their visible, left hand on the touch screen mounted directly above the robot handle where they thought their trained hand had ended its movement under the arc.

Crucially, there were two variations of this task. First, in the ‘active’ localization task participants generated the movement themselves, as they could freely move their unseen right hand from the home position to any point on the arc (Fig 2a, dark blue boxes). Second, there was a ‘passive’ localization task where the robot moved the participants’ hand out and back (Fig 2a, light blue boxes), to the same locations the participants moved to in the preceding ‘active’ localization task in a shuffled order. Hence, active localization was always done first. In active localization, participants had access to both afferent, proprioceptive information as well as an efference-based prediction of sensory consequences, but in passive localization, only proprioception should have been available. The active and passive localization task each consisted of 25 trials, and each of the tasks was done a total of four times; twice after aligned and twice after rotated training. From these tasks, the localization responses were analyzed, i.e. where participants indicated the endpoint of their hand movement was, in particular the angles of the indicated points relative to the home position in comparison to the true reach endpoints’ angle.

### Classic training

The paradigm described above was an exact replica of a paradigm we used earlier [9] with three exceptions. First, we used exposure training here, instead of the standard reach training with volitional movements, which we called ‘classic’ training. Second, all localization was delayed until the right hand has returned to the home position in this study, whereas the previous study also included a second set of “online” localization tasks, where touch screen responses were collected when the unseen, right hand was still at the furthest point of the movement. Instead of both delayed and online localization we had two repetitions of each delayed localization task here. Third, we added cursor blink detection to the training tasks, to make sure participants attended to the task. In the present study we compared changes in localization responses and no-cursor reach endpoints after exposure training with changes in the same measures after classic visuomotor adaptation training, as collected earlier.

### Analyses

Prior to any analyses, both the localization responses and the no-cursor reach data were visually inspected and trials where the participants did not follow task instruction were removed (e.g. several movements back and forth, or a touch-screen response on the home position, instead of on the white arc). From the localization responses 1024 / 16800 (^~^6.1%) trials were removed from the exposure group’s data, and 220 / 8400 (^~^2.6%) from the classic group’s data. From the no-cursor reaches, 2 / 3984 (^~^0.05%) were removed from the exposure group’s data, and 1 / 3969 (^~^.025%) was removed from the classic group’s data.

### Localization

We primarily wanted to test if hand localization shifts after exposure training; whether or not there were differences between active and passive localization shifts; and finally compare localization shifts after exposure training to those after classic visuomotor training.

There were some idiosyncratic differences in performing localization (e.g. systematic under- or overshoot of the arc by some participants). To counter this, we fitted a circle with a 10 cm radius to the touch screen responses of each participant (using a least-squared errors algorithm) and the offset of this circle’s centre was subtracted from all response coordinates, so that all responses fell close to the arc. Localization responses were taken as the (signed) angular difference between vectors through the home position and both the actual hand position as well as the location indicated on the touch screen. For each localization response, we retain the angle of the actual reach endpoint, as well as the signed angular error of the localization response. Since participants freely chose their movement directions, we interpolated angular localization errors at the same angles used for the no-cursor reaches (15°, 25°, 35°, 45°, 55°, 65° and 75°), but only if that location fell within the range of the data (i.e. we did not extrapolate). For interpolation we used a smoothed spline that was fit to every participant’s response errors in each of the four localization tasks (aligned vs. rotated and active vs. passive). This way localization responses could be compared across participants despite the freely chosen reach directions. At the 15° location 7/21 participants didn’t have an estimate in one or more of the four localization tasks (in the “classic” data it was also 7/21). While that data was shown in the figures, we did not use it for analysis.

First we compared localization responses shifted following rotated exposure training to those following aligned, and confirmed that training shifted localization. We then tested if the shift in localization responses is different for active and passive localization, and we ran analyses comparing localization after exposure training with localization after “classic” training. Finally, we test if the variance of active localization responses is lower than the variance of passive localization responses.

### Reach aftereffects

For our secondary question, we wanted to assess if any reach adaptation had occurred after exposure training, so we analyzed reach endpoint errors in no-cursor trials. Reach endpoint errors were the (signed) angular difference between a vector from the home position to reach endpoint and a vector from the home position to the target. We obtained reach aftereffects by subtracting median reach endpoint errors for each of the 7 targets after aligned training from those after rotated training. No-cursor endpoint errors were analyzed to test if participants adapted the direction of their reaching movements after rotated exposure training, which would indicate that recalibrated proprioception contributes to motor changes. We also tested if any such change decayed, i.e. if it was the same immediately after exposure training, or when a localization task was done in between exposure training and no-cursor reaches. Finally, we compare reach aftereffects between the exposure and classic training groups.

### Generalization maxima

We noticed that the generalization curves after exposure and classic training looked different, and the planned analyses indeed showed interactions between hand angle and training paradigm. Specifically, the maxima after classic training are very close to the target, as expected, but for exposure training, the maxima seemed shifted (see Results). Since this was potentially interesting, we added two purely exploratory (i.e. unplanned) analyses of the location of the maxima in the workspace. The same analysis was done for active localization and reach aftereffects, comparing generalization after exposure and classic training. Note that these analyses did not answer our main questions.

Pre-processing and analyses were done in R 3.4.4 [28] using lme4, lmerTest and various other packages. There is some systematically missing localization data, which means we can’t run ANOVA’s on the full data set. We also can’t reliably extrapolate or impute beyond the range of responses, and removing hand locations (4/7) or participants (7/21) would remove too much data, decreasing power. Hence we decided not to use regular ANOVAs but linear mixed effects models instead, as these are robust against missing data. The output was converted to more readable ANOVA-like output, using a Satterthwaite approximation [29]. Highly similar results were obtained with a Chi-square approximation. Data, scripts and a notebook with analyses have been made available on the Open Science Framework [26] (https://osf.io/zfdth).

In short, this experiment allowed us to test how mere exposure to a visual-proprioceptive discrepancy changed both reach aftereffects and hand localization responses, and compare them with those obtained after more common, “classic” training.

## Results

In this study we intend to further elucidate the relative contributions of (updated) predicted sensory consequences and (recalibrated) proprioception to training-induced shifts in hand localization. We can parcel out these contributions by measuring hand localization after both robot-generated (afferent signals only) and self-generated movements (both afferent and efferent signals). Finally, all analyses are performed with the data from the current experiment and those obtained in an earlier study that uses an identical paradigm, but training with self-generated movements, or “classic” training.

### Localization

Here we test our hypothesis that exposure training with a visual-proprioceptive discrepancy does not lead to changes in predicted sensory consequences, which predicts equal shifts in active and passive localization. First, we can see that the difference between rotated and aligned localization responses is different from zero, after exposure training (Fig 3a), so that it seems there are *shifts* in localization that we can compare between tasks and training paradigms. To verify there is a shift in localization responses, we fit an LME model to the localization errors throughout the workspace using *session* (aligned or rotated), *movement type* (active and passive) and *hand angle* (25°, 35°, 45°, 55°, 65° and 75°), and all interactions as fixed effects and *participant* as random effect. There is an effect of *session* (F(1,450.5)=155.8; p<0.001), showing that training with exposure to a visual-proprioceptive discrepancy leads to a shift in hand localization. There is also an effect of *hand angle* (F(5,450.6)=6.54; p<.001) and an interaction between *hand angle* and *session* (F(5,450.3)=8.25; p<.001), but no other effects (all p>.60). Since exposure training led to a shift in localization responses, we use the difference between hand localization after rotated training and after aligned training as a measure of localization shift.

**Fig 3.**
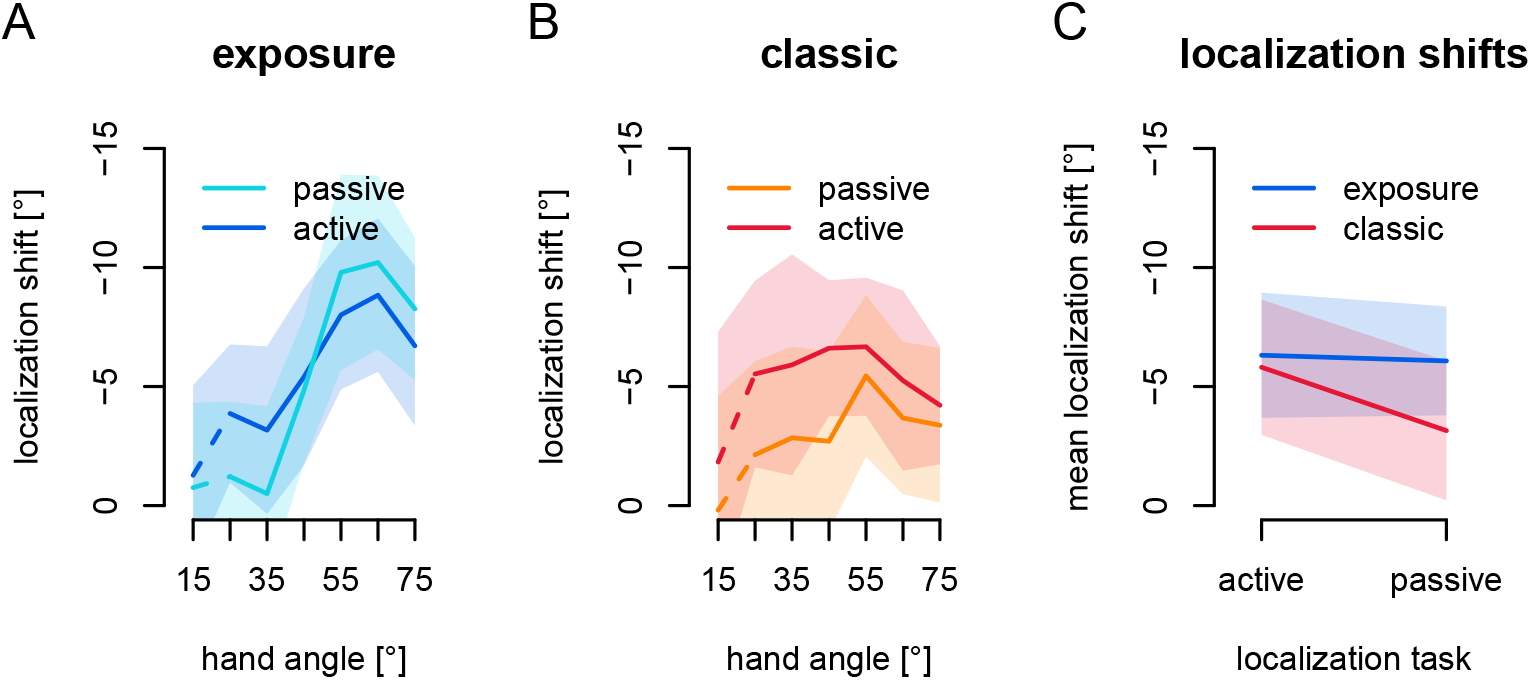
Hand Localization. The shifts of the angles of touchscreen responses in all variations of the localization task, using spline-interpolated estimates for hand angles matching the reach targets in the no-cursor reach tasks. Shaded areas denote 95% confidence intervals. **a)** Localization shifts after exposure training. *Dark blue*: active localization shifts, *Light blue*: passive localization shifts. **b)** Localization shifts after classic training. *Dark red*: active localization shifts, *Orange*: passive localization shifts. The dashed line segments illustrate that the 15° data is not used for statistical analyses (see Methods). **c)** The overall localization shifts for active and passive localization after both exposure and classic training. *Blue*: localization shifts after exposure training, *Red*: localization shifts after classic training.

Since the difference between active and passive localization stems only from the presence or absence of predicted sensory consequences, there should be no difference between the two if predicted sensory consequences are not updated by exposure training. At first glance, it seems there might be a difference between active and passive localization shifts in exposure training (see Fig 3a), although it is smaller than in classic training (Fig 3b). Also noteworthy is that the overall magnitude of the localization shifts seems higher after exposure training as compared to classic (Fig 3c). This is unexpected, but we do find a similar pattern in other work [23].

If, contrary to our hypothesis, this shift in localization after exposure training partly reflects predicted sensory consequences, then shifts in active localization, that rely on both (recalibrated) proprioception and (updated) predictions should be different from shifts in passive localization that only rely on (recalibrated) proprioception. We fit a linear mixed effects model (LME) to the change in localization using *movement type* (active or passive localization) and *hand angle*, as well as their interaction as fixed effects and *participant* as random effect. As we expected, there is no effect of *movement type* (F(1,211.8)=0.07; p=0.79). There is an effect of *hand angle* (F(5,212.2)=10.8; p<0.001), but no interaction between *hand angle* and *movement type* (F(5,211.8)=1.23, p=.29). The lack of an effect of *movement type* means we don’t have evidence that predicted sensory consequences contribute to localization in this paradigm.

Active localization shifts are larger than passive localization shifts after classic training (Fig 3b). In order to compare hand localization shifts after exposure training with those after classic training [9], we fit an LME model to localization shifts using *training type* (exposure vs. classic), *movement type* (active vs. passive) and *hand angle* and all interactions as fixed effects and *participant* as random effect, and expect an interaction between *training type* and *movement type*. There is a main effect of *movement type* (F(1,422.0)=6.22; p=.013) and of *hand angle* (F(5,422.7)=8.19; p<.001), as well as an interaction between *training type* and *hand angle* (F(5,422.7)=4.54; p<.001) and between *training type* and *movement type* (F(1,422.0)=4.48, p=.035), but there is no main effect of *training type* (F(1,39.1)=0.92, p=.34) and no other effects (all p>.14). These results suggest that the magnitude of the shifts in localization are comparable between classic and exposure training (no main effect of *training type*), but that the pattern of generalization is different (interaction between *training type* and *hand angle*). Importantly, the interaction between *training type* and *movement type* is in line with our main prediction that the difference between active and passive localization shifts should be different in the two training types.

To further address our main question, we will consider the interaction between *training type* (exposure vs. classic) and *movement type* (active vs. passive) we report above (see also Fig 3c). The overall larger shifts after exposure training could be due to inherent differences between the groups, or because either exposure training magnifies proprioceptive recalibration, or because the sensory prediction error based learning present in classic training dampens other mechanisms [27], such as proprioceptive recalibration. However, since there is no difference between active and passive localization shifts after exposure training alone, the interaction between *training type* and *movement type* should be caused by an effect of *movement type* on the localization shifts after classic training, as we found previously [9]. This strongly suggests that shifts in hand localization after exposure training indeed rely on recalibrated proprioception alone, while after classic training, there also is a contribution of predicted sensory consequences to active localization.

### Localization variance

In active movements there could be more information available to localize the hand, which could lead to more precise responses. To test this, we calculate the variance of localization responses relative to a smoothed spline expressing the bias of localization responses in each participant across the workspace. A paired-sample t-test shows that the variance of active and passive localization are not different from each other in the aligned session (t(20)=1.178, p=.253) as well as the rotated session (t(20)=1.066, p=.299) in the exposure training group. This confirms earlier studies where we also do not find that active localization is more precise than passive localization [6, 9, 30].

### Reach aftereffects

Apart from our main question on the contribution of proprioception and prediction to hand localization, we want to see if rotated exposure training has any effect on open-loop reaches and if these persisted. We quantify whether participants adapt their reach directions by assessing their reach errors in no-cursor reach trials after aligned and rotated exposure training. In Fig 4, the changes in no-cursor endpoint errors, or reach aftereffects, appear to be well over 5°. First, to confirm that exposure training affects open-loop reach direction, we fit a linear mixed effects model (LME) to reach endpoint error using *session* (aligned; all blocks, or rotated; only the first block immediately after training) and *target* (15°, 25°, 35°, 45°, 55°, 65° and 75°), as well as their interactions as fixed effects and *participant* as random effect. There is an effect of *session* (F(1,260)=93.81, p<.001), but there is no effect of *target* (F(6,260)=1.07, p=.37) and no interaction (F(6,260)=1.00, p=.42). Since there is an effect of *session*, we can say there are reach aftereffects, and now take the differences in reach endpoint errors between the rotated and aligned session for every participant and target as “reach aftereffects”, and use those for all further analyses. Importantly, the effect of *session*; i.e. open-loop reaches after rotated exposure training are different from those after aligned training, also indicates that exposure training evokes motor changes usually associated with other adaptation mechanisms.

**Fig 4.**
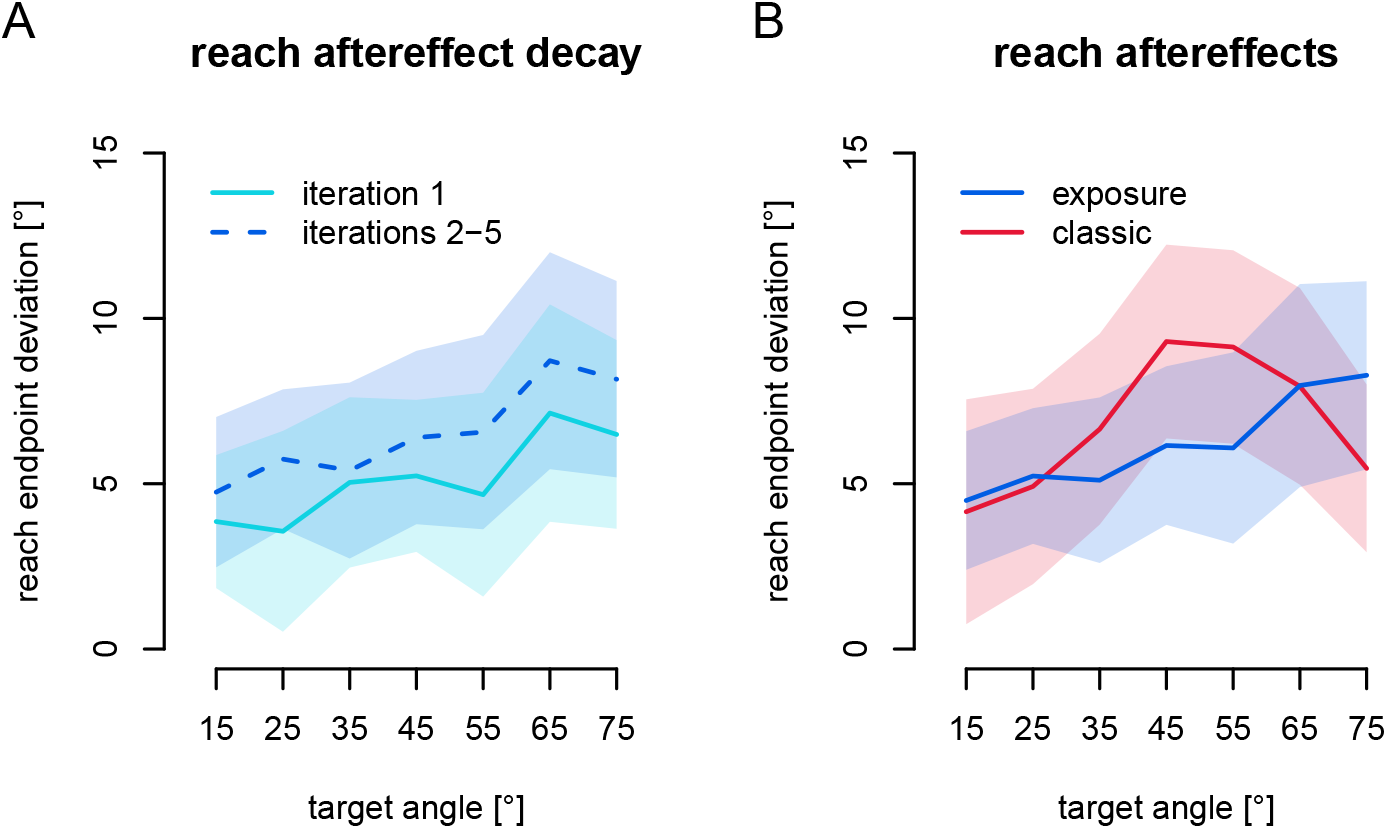
Reach aftereffects. Changes of the angle of reach endpoints in the no-cursor tasks after rotated training. Shaded areas denote 95% confidence intervals. **a)** Reach aftereffects at different time points in the rotated session. *Light blue*: first no-cursor task in the rotated session (immediately following training), *Dark blue, dashed* line: the other four repetitions of the task (with localization in between training and no-cursor tasks). **b)** Reach aftereffects after classic and exposure training. *Blue*: exposure training, *Red*: classic training.

To see if reach aftereffects decay during the localization tasks, we compare reach aftereffects in the initial no-cursor block, that immediately follow training, with those in the later blocks that follow a localization task (Fig 4a). We fit an LME model to the reach aftereffects with *iteration* (initial vs. later no-cursor blocks) and *target* (as above) as well as their interaction as fixed effects and participant as random effect. There is no effect of *iteration* (F(1,260)=2.72, p=0.10). There is an effect of *target* (F(6,260)=6.29, p<.001) but no interaction (F(6,260)=0.58, p=0.74). In sum, reach aftereffects are not appreciably different right after training or after the localization tasks. In other words, there is likely no noticeable decay of reach aftereffects during the localization tasks, so that we can collapse the data across iterations.

Next we compare the reach aftereffects after classic training with those after exposure training (Fig 4b). It appears as if there is little overall difference in magnitude, but there might be a shift of the generalization curve. We fit an LME model to reach aftereffects with *training type* (classic vs. exposure), *target* (as above) as well as their interaction as fixed effects and *participant* as random effect. Contrary to our expectation, and earlier findings [13], there is no main effect of *training type* (F(1,40)=0.11, p=.74), indicating approximately equal overall magnitude of reach aftereffects after the two training types. There is an effect of *target* (F(6,240)=8.36, p<.001), indicating that reach aftereffects exhibit some form of a generalization curve. There is also an interaction between *training type* and *target* (F(6,240)=2.27, p=.038), indicating these generalization curves are different after the two training types. To summarize, we find that exposure training elicits substantial and persistent motor changes and we don’t find a difference in the magnitude of reach aftereffects between those after classic and exposure training.

### Generalization maxima

While not our main question, we will add exploratory analyses on the potentially different generalization patterns of localization shifts and reach aftereffects after classic and exposure training (Fig 5). The LME models above indicate no difference in overall amplitude of localization shifts or reach aftereffects between the groups, so the interaction between training type and hand angle (or target for reach aftereffects) might stem from a shifted generalization curve after exposure training, in contrast to classic training where the maximum is close to the trained target [31]. Using the active localization shifts only (which are larger, and should arguably be more similar to reach aftereffects, because they include predicted sensory consequences), we bootstrap a 95% confidence interval for the maximum localization shift, by resampling participants within each group 100,000 times and taking the centre of a normal curve fit. This procedure indicates that after classic training, the maximum localization shift is at 48.3° (95% confidence: 36.9° - 57.8°; red area in Fig 5a), and after exposure training the maximum localization shift is at 62.5° (95% confidence: 55.4° - 70.8°; blue area in Fig 5a). This means that the maximum effect of classic training on localization, occurs at a different point in the workspace than that of exposure training.

**Fig 5.**
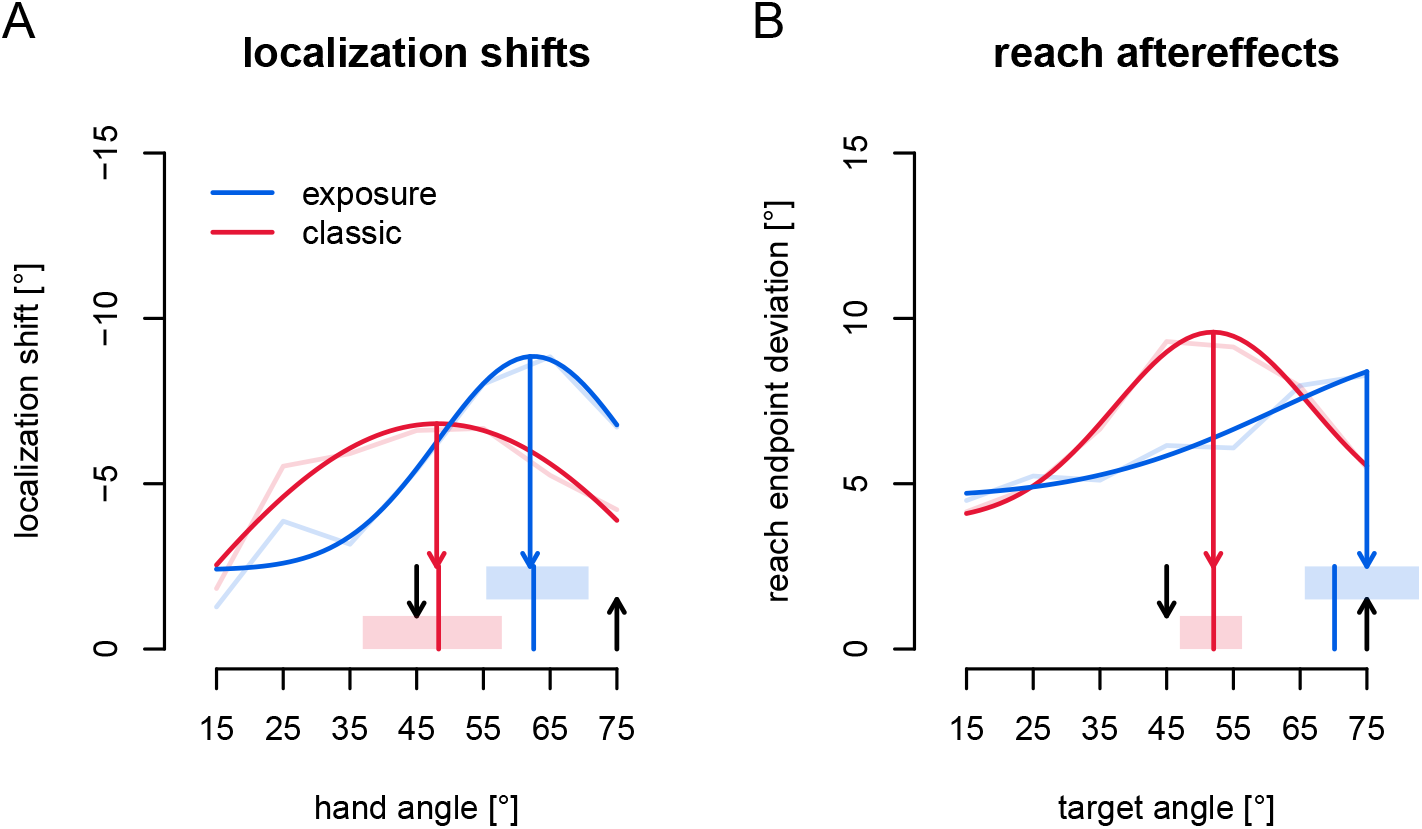
Generalization maxima. Locations of generalization maxima in the workspace for active localization and reach aftereffects. *Blue*: Exposure training, and *red*: classic training. The average data is shown as faint lines, and the curves are Gaussians fitted to all the data. Shaded areas at the bottom mark the 95% confidence intervals for the location of the maximum of each curve (vertical red and blue lines in shaded areas indicate the median; colored arrows pointing downward from the curves indicate the maxima of the curves fitted to all data). Downward black arrows: visual trained target at 45°. Upward black arrows: hand location during training at 75°. **a)** Generalization maxima of active localization shifts. **b)** Generalization maxima of reach aftereffects.

In Fig 5b we can observe a similar pattern for reach aftereffects. Using the same bootstrapping procedure, we find that the maximum of the generalization curve of reach aftereffects after exposure training is at 70.2°, with a 95% confidence interval ranging from 65.7° to 153.5°. This very high upper bound can be explained by the fact that our experiment didn’t sample the full distribution (nor was it meant to), so that a curve with a peak at such a location could be an accurate fit of the data. However, we can at least take the lower bound as the minimum location in the workspace where reach aftereffects maximally generalize, as repeated bootstrapping procedures all converge on values around 65°. In comparison, the generalization of reach aftereffects after classic training, peaks at 52.0°, with a 95% confidence interval spanning 47.0° to 56.3°. This means that the generalization curves for reach aftereffects after exposure and classic training have maxima at different directions relative to the trained direction.

The LME models for localization shifts and reach aftereffects indicate a different generalization curve after exposure and classic training, and this can be explained by a different position of the generalization curves after the two types of training. The unrealistically high upper bound for the confidence interval of the peak of the generalization curve of reach aftereffects after exposure training, indicates that we should sample a wider part of the workspace in future experiments to get more accurate fits. However, the different generalization functions after classic and exposure training indicate that the mechanisms of adaptation that are employed by participants in each of the training paradigm are substantially different, as we have found before [11], even when both affect hand localization and open-loop reach directions. In classic training, sensory prediction errors mainly drive adaptation [27], and these are more or less anchored to the visual target at 45° [31, 32]. In exposure training, cross-sensory discrepancy drives adaptation, such that the experienced afferent signals from the trained hand position (75°) are newly associated with the rotated visual feedback. Most likely, the different error signals underlying each type of training are the source of the differences in generalization. An open question is how these two generalization functions interact in behavior.

The differences between the generalization curves for both reach aftereffects and hand localization shifts after both classic and exposure training are not well understood. Perhaps the error-signals underlying both may provide a hypothesis as the basis for future work. On the one hand, in classic training, what gets updated is the movement that should be used to reach a certain visual target using a visual cursor. Hence the reach aftereffects, and perhaps hand localization shifts should be strongest close to the visual target [31]. On the other hand, in exposure training, people do not have to make any movements, but the proprioceptive feedback they experience is associated with different visual feedback. Hence the hand localization shifts, and perhaps reach aftereffects should be strongest close to the experienced hand position. This hypothesis matches the patterns we observe here, but requires experimental verification.

Regardless of the differences in generalization maxima, exposure training leads to substantial shifts in hand localization that are not different for active or passive localization, while movement type does have an effect on localization shift after classic training. Exposure training also causes persistent reach aftereffects that are not different in magnitude to those found with classic training.

## Discussion

Accurate information on where our limbs are is important for planning and evaluating movements, and can be estimated through predicted sensory consequences, as well as visual and proprioceptive feedback. As in a previous study [9] here we quantify the contributions of predicted sensory consequences and proprioceptive recalibration to where we localize our hand after training with altered visual feedback of the hand. In classical adaptation paradigms, both predictions are updated and proprioception is recalibrated. Predictions are updated when they don’t match actual sensory consequences, and proprioception is recalibrated when it doesn’t match visual feedback. In this study we use “exposure” training, where the participants do not have volitional control of their movements. By design, this should eliminate efference copies and prevent updating predicted consequences of movements, but since the proprioceptive and visual feedback is the same, exposure training still allows proprioceptive recalibration. Before and after training, participants localize their hand, both after “active,” self-generated movements that allow using predicted sensory consequences, and after “passive,” robot-generated movements that only allow using proprioception. We calculate the training-induced shift in both types of localization given the same actual hand position. After classical training we previously reported larger shifts in active localization as compared to passive [9]. As we expected, after exposure training there are substantial shifts in localization, but no differences between active and passive localization, indicating that predictions are not updated after exposure training. Somewhat unexpectedly, localization shifts after exposure training are slightly larger than after classic training. Furthermore, we find that exposure training evokes substantial and robust reach aftereffects, indicating that recalibrated proprioception is used to plan movements.

Our lab previously investigated proprioceptive recalibration and reach aftereffects following visuomotor adaptation with classic training and matched exposure training, however without comparing active with passive localization. In those studies we usually find that proprioceptive recalibration is of similar magnitude in both training paradigms, but unlike here, reach aftereffects are usually much larger with classic training [13, 18, 23–25]. In one study [23], we observed that localization shifts after exposure training are larger than after classic training – like here. Perhaps focusing on the sensory information increases proprioceptive recalibration in exposure training, or engaging in sensory prediction error based learning reduces the need to engage other adaptive mechanisms, such as proprioceptive recalibration. And while proprioceptive recalibration and reach aftereffects do proportionally increase with gradual increases in rotation size for classical training, they do not for exposure training [25]. The similar magnitude of proprioceptive recalibration and reach aftereffects following exposure training, but not classical training, suggest that sensory recalibration is driving this modest change in movements, with additional motor changes after classic training through updates of internal models. The effect of exposure training on movements is also demonstrated by savings and interference from exposure training carrying over to subsequent classic training [33] and transfer of exposure training effects from one hand to the other [34]. In the current study, we further demonstrate that exposure training affects both movements and proprioception, and verify that it likely has little effect on predictive estimates.

Results similar to what we find here were reported in a study by Cameron and colleagues [19], using gain modulation of visual feedback of single-joint hand movements around the elbow. Their within-subjects experiment includes both training with volitional movements as well as with machine-generated movements and also tests perception of movements that were either passive or active. They too find a robust change in passive perception of hand movement (using a different measure), and these changes do not differ between the two types of training. Similarly, they find shifts in what we might call “active localization,” although the task is different, after both training types. These shifts are larger after classic training as compared to exposure training. They also find that exposure to a visuo-proprioceptive discrepancy leads to reach aftereffects, although these are smaller than those produced following “classical” training with altered visual gain. Both our findings, and those of Cameron et al. [19] indicate that updating predicted sensory consequences requires volitionally controlled movements that lead to sensory prediction errors, while proprioception recalibrates equally in both types of training, and that recalibrated proprioception affects open-loop reaches. Our combined results suggest that updates in predicted sensory consequences only provide a partial explanation for visuomotor adaptation.

Two related concerns about exposure training and passive localization are that first, the movements might not be not fully passive, so that efference-based predictions are still generated, and second, that predicted sensory consequences could be generated through another route, e.g., based on afferent signals. Cameron et al. [19] measured muscle activity (EMG) during passive‘ movements and found no difference with stationary baseline muscle activity. This implies that no movements are generated in any passive condition or they are subthreshold, so that efference-based predictions are at worst minimal. The brain areas generating predicted sensory consequences could also rely on afferent signals. However, such afferent signals are present in both active and passive movements, and if they would result in the same predictions, there would be no difference between active and passive localization after classic training, but there is. Hence, while we can not fully exclude any predictive signals in passive localization or exposure training, our data shows that any residual predictive signals in the passive movements we used are qualitatively very different from normal efference-based predicted sensory consequences.

Apart from these concerns, it might be that active localization is more precise than passive localization, as it is based on more information. However, here we confirm previous findings [6, 35] that show this is not the case, and recently replicated this in a much larger dataset [36]. This sets limits for the kinds of mechanism we can hypothesize are used by the brain to combine efferent with afferent signals, as well as perhaps the nature of the afferent signals used in localization. Assuming that predictions were not updated in exposure training, a maximum likelihood estimate (MLE) or “optimal integration” [37] would predict that active localization should shift less than passive localization after exposure training. But this is not the case in our findings (although it is the case for Cameron et al., [19]). Taken together, this suggests these different sources of information about unseen hand location are probably not optimally integrated. It also suggests that the afferent signals in active localization are not more informative than afferent signals in passive localization. While localizing the unseen hand is less precise than locating (pointing to) a remembered visual target or a seen and felt hand location, we find that these bimodal estimates are rarely integrated optimally [35, 38, 39], although others have [40]. A more recent study [41] has also shed doubt on whether “optimal” or “Bayesian” integration is used for locating the hand with two afferent signals. Analogously, here we again can’t find evidence that afferent and efferent information combine as a maximum likelihood estimate.

It seems clear that the cerebellum plays a role in motor learning as it appears to compute predicted sensory consequences, i.e. it implements a forward model [42–44]. People with cerebellar damage do worse on motor learning tasks [45–48], and show decreased shifts in hand localization tasks following motor learning [5, 7]. This highlights that the cerebellum, and likely predicted sensory consequences, are important for motor learning, but does not explain the remaining shifts in hand localization. We previously found that proprioceptive recalibration is intact in people with mild cerebellar ataxia and that it is similar following exposure and classical training with a gradually introduced cursor rotation [18]. The remaining changes in hand localization found in cerebellar patients can be attributed to recalibrated proprioception which should be intact [18]. Analogously, here we show that in a paradigm that stops updates of predictions of sensory consequences, as supposedly in people with cerebellar damage, we still see substantial shifts in localization. Again, the remaining localization shifts can be explained if, along with predictions, the human brain uses afferent signals: recalibrated proprioceptive estimates, to localize the hand.

## Conclusion

To sum up, after a training paradigm designed to prevent updating of predicted sensory consequences but allowing recalibration of proprioception, we find substantial changes in where people localize their hand. This means that recalibrated proprioceptive estimates can explain shifts in hand localization. Since our participants also changed the direction of open-loop reaches, recalibrated proprioception seems to guide motor planning. This study confirms that sensory prediction error based learning and proprioceptive recalibration are different mechanisms that separately contribute to motor adaptation.

## Acknowledgements

This work was supported by a Canadian Network for Research and Innovation in Machining Technology NSERC Operating grant (DYPH) and the German Research Foundation (DFG) under grant no. HA 6861/2-1 (BMtH). The funders had no role in study design, data collection and analysis, decision to publish, or preparation of the manuscript. We thank Shanaathanan Modchalingam for assistance in collecting the data.

